# Challenges in the calibration of large-scale ordinary differential equation models

**DOI:** 10.1101/690222

**Authors:** Eva-Maria Kapfer, Paul Stapor, Jan Hasenauer

## Abstract

Mathematical models based on ordinary differential equations have been employed with great success to study complex biological systems. With soaring data availability, more and more models of increasing size are being developed. When working with these large-scale models, several challenges arise, such as high computation times or poor identifiability of model parameters. In this work, we review and illustrate the most common challenges using a published model of cellular metabolism. We summarize currently available methods to deal with some of these challenges while focusing on reproducibility and reusability of models, efficient and robust model simulation and parameter estimation.

## 1 Introduction

Understanding complex biological processes is the key goal of of systems biology (Kitano, 2002). Mathematical models, in particular based on ordinary differential equations (ODEs), have been playing an important role in this context (Kholodenko et al., 1999; Kollman et al., 2005). Some processes can be explained with comparably simple models (Becker et al., 2010). Others, such as the interplay of multiple pathways in complex diseases or the functioning of whole cells, need large models to be described (Hass et al., 2017; Karr et al., 2012; Swainston et al., 2016). Due to the advances in experimental and computational methods, many large-scale models have been developed in recent years (Froehlich et al., 2018; Khodayari and Maranas, 2017; Mazein et al., 2018). As the data acquisition speed continuously increases (Barretina et al., 2012; Li et al., 2017) and more sophisticated tools for model development appear (Gyori et al., 2017), this trend is likely to persist.

In the past years, computational tools for the creation and calibration of ODE models have been developed and have shown to perform well for small- to medium-scale models (Henriques et al., 2017; Hoops et al., 2006; Raue et al., 2015). Efficient methods have been developed for model calibration and uncertainty analysis (Ballnus et al., 2017; Raue et al., 2013; Villaverde et al., 2018). This has facilitated the process of model development and calibration. However, many of these methods do not scale well to large models sizes with thousands of state variables or parameters. Such limitations have been observed in recent studies using large-scale models (Froehlich et al., 2018). During the establishment of a comprehensive benchmark collection (Hass et al., 2019), we experienced that even the reimplementation of existing models and the reproduction of published results becomes increasingly difficult with model size.

In this review, we summarize some of the most common challenges in the (re-)use, simulation and parameter estimation for large-scale ODE models, present current approaches to deal with them and use a recently published E. coli metabolic model as application example. We do not cover the topics of model construction or data generation. Every subsection in the main part of this work first discusses the general case, covers the application example in the penultimate paragraph and then concludes with a short summary.

## 2 Application example and notation

Throughout this manuscript, we use the ODE model k-ecoli457 by Khodayari and Maranas (2017) as an application example. The model describes the dynamics of 3003 biochemical species (metabolites, enzymes and enzyme-metabolite complexes) using 5239 elementary reactions. The reactions follow mass action kinetics, yielding 5239 kinetic parameters. Following the work of Khodayari and Maranas (2017), we used 31 steady state flux distributions of E. coli including 25 flux distributions grown under aerobic conditions with glucose as carbon substrate, three flux distributions grown under aerobic conditions with pyruvate as carbon substrate, two flux distributions grown under anaerobic conditions with glucose as carbon substrate, and one flux distribution grown under aerobic conditions with acetate as carbon substrate. Model and experimental data were extracted from the supplementary information of the publication and comprise 12 to 46 measurements for each growth condition, adding up to a total of 1047 data points. Our aim was to use this model as a case study for comparing current tools and methods on it. As this turned out to be more challenging than expected, we decided to summarize the points which we imposed the hardest challenges to us.

We consider ODE systems of the form

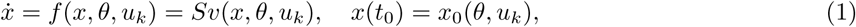

in which *x* denotes the vector of state variables, *x*_0_ its initial values, *u*_*k*_ a vector of external inputs for an experimental condition *k* = 1, …, *n*_*e*_, *S* the stoichiometric matrix, *v*(*x, θ, u*_*k*_) the reaction flux vector, and 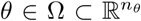 the unknown parameters of the model, which are restricted to a biologically plausible region 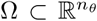. The input vector *u*_*k*_ is assumed to capture all differences between experimental conditions. A steady state of the model is denoted by

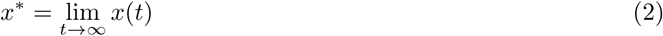

with

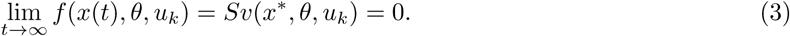

Following the publication of Khodayari and Maranas (2017), we assumed a subset of the reaction flux vector as observable quantities, which we compared with the measurement data. As measurements are noise corrupted, we assumed an additive normally distributed noise model with fixed standard deviation. Based on this noise model, we used the negative log-likelihood of observing the data given the parameters *θ* as objective function.

## 3 Challenges and possible approaches

### 3.1 Model formulation and reusability

The first challenge faced when studying large-scale ODE models is to actually implement them. Publications sometimes provide the underlying reaction networks, sometimes the ODEs and their initial conditions, maybe with executable code, or only (incomplete) descriptions in form of written text. Depending on the form of publication, reimplementing an ODE model with thousands of reactions may take days or weeks and can be extremely error prone.

While publishing an ODE model as right hand side with initial conditions and executable code ensures reproducibility -i.e., the possibility to reproduce the shown results -this does not hold true for reusability -i.e., the possibility to use this model for other purposes than just reproducing known results. A good and widely used way to ensure reusability is publishing the model as reaction network with its kinetics in one of the XML-based formats SBML (Hucka et al., 2003) or CellML (Miller et al., 2010). Most of the current toolboxes for model simulation and parameter estimation support at least SBML and allow for automated import and translation to an ODE model. Depending on the model size, this takes seconds to minutes. Hence, formulating and publishing a model in SBML will substantially facilitate its reuse and increase its impact.

In our considered application example, the stoichiometric matrix, the reaction fluxes and the ODE were provided as MATLAB code. Model parameters and state variables were documented in separate XLS-sheets. This made it easy to run the simulations from MATLAB directly, but very challenging to reuse it for other tasks. We tried to recreate an SBML file from the right hand side using MOCCASIN (Gomez et al., 2016), which however failed due to the model size. Hence, we had to reimplement the right hand side, which took us many days until the simulations of all experimental conditions were running correctly. Given an SBML file, running the corresponding import function would have taken less than an hour.

In summary, the use of existing standards for model formulation is highly beneficial. This becomes even more important for large-scale models, as the reimplementation effort increases with model size. Publishing an SBML model on respective databases such as BioModels (Li et al., 2010) or JWS online (Waltemath et al., 2017) may even further encourage its reuse and increase its impact.

### 3.2 Availability of measurement data and link to the model

The next challenge, which concerns measurement data, is two-fold. As firstly, large-scale models usually possess many unknown parameters, large data sets are required to ensure that the model calibration problem is not underdetermined. At least as many data points as model parameters should be used to ensure the theoretical possibility of determining the model parameters. In practice however, substantially more data points may be needed. Hence, if possible, already published data may be used or prior information about reaction kinetics implemented, e.g. from databases like BRENDA (Schomburg et al., 2013) or SABIO-RK (Wittig et al., 2012). In our application example, about 1000 data points were used for more than 5000 parameters. Consequently, most of the model parameters could not be determined, which may have substantial drawbacks on the quality of model predictions.

The second challenge consists in linking the existing experimental data to the model simulations. Meta-data, e.g. experimental conditions, are required and data processing, such as data normalization, must be taken into account by introducing scaling factors or measurement offsets, when observable quantities of the system should be modelled appropriately. Often, such crucial information is missing in public data sets and the data mapping gets increasingly harder the larger the data set is.

Ideally, this challenge would be addressed by using standard formats for observable quantities of the model, formulation of measurement data and combining this information for parameter estimation. Unfortunately, such standards are currently missing. The XML-based format SBRML (Dada et al., 2010) was designed for this tasks, but was not adopted by the scientific community. SED-ML (Waltemath et al., 2011) is another standard for formulating observable quantities and experimental description. However, also this standard is not widely in use and currently only a small number of toolboxes provide an interface. Moreover, SED-ML is missing some specifications which would make it applicable for the direct use in parameter estimation, such as the possibility to indicate noise models, to implement an objective function, or employ parameter priors. Therefore, most parameter estimation toolboxes have their own specific formats, which makes it hard to switch between the toolboxes for different tasks. Community efforts, such as PEtab (Weindl et al., 2019), exist to tackle this problem, but are not yet applicable to a wide range of models and data types.

In the case of our application example, we reimplemented the 31 experimental conditions, ran simulations, collected the experimental data, and compared them to the model output. Despite a good documentation from the authors, this took – also due to the high computation time – several days. During this process, we found two flipped minus signs in the formulation of the observed metabolic fluxes (as confirmed by the authors). The authors had fixed them correctly in their parameter estimation code, but had described it neither in the publication nor in the provided code. This illustrates that handcrafted solutions are likely to produce errors and should therefore be avoided.

Currently, SED-ML is the most promising candidate for providing experimental conditions and observable quantities in a simulation-ready format and should therefore be considered an option. However, standards which allow the formulation of a parameter estimation problem are missing and currently being developed. Thus, a detailed documentation of the parameter estimation problem is probably the best way to ensure an easy reproduction and reuse of published results.

### 3.3 Efficient simulation of ODE models

A key challenge is achieving an efficient and reliable forward simulation of the model. During model calibration or uncertainty analysis, this has to be done thousands or even millions of times. As in many projects, computation time is the limiting factor, an efficient model simulation is urgently needed. Due to their size, large-scale models are often more prone to numerical artifacts or simulation failures. Because of these numerical problems, measures must be taken to ensure not only an efficient but also a robust model simulation.

The first step consists in choosing the most appropriate approach for forward simulation: If time series data is to be simulated, the ODE must be solved, which is typically done by numerical integration with implicit solvers. For the study of steady state data, the model can either be simulated to a late time point until some norm of the right hand side undergoes a previously chosen threshold, or root finding algorithms, such as Newton’s method, can be used.

An approach to improve the efficiency of implementations is the use of compiled languages. Mature toolboxes for ODE solving or root finding are available (Hindmarsh et al., 2005; Zhang and Sandu, 2011) and should be considered, unless a custom implementation is unavoidable. Often, those toolboxes can be interfaced from high-level languages, such as R (Soetaert et al., 2010) or MATLAB and python (Fröhlich et al., 2017b), to facilitate their usage. This may decrease computation time by orders of magnitude.

Improving robustness is often model specific and highly challenging. Generally, the output of solver diagnostics, will often be helpful to spot numerical issues in the case of simulation failures. During ODE integration, entries in the state vector, such as concentrations, which are physically required to be positive, may become negative if inappropriate error tolerances are used. In these cases, it may help to enforce positivity of the state vector or slightly adapt the reaction kinetics, if such an adapted model is still considered appropriate. If Newton’s method is used, it may happen that the Jacobian of the right hand side is singular and can not be factorized. To avoid singularity of the Jacobian matrix as far as possible, the system size should be reduced by exploiting conservation laws, which is done automatically in some of the common toolboxes (e.g., COPASI) or shown in (Vallabhajosyula et al., 2006).

We simulated the application example using the ODE solver toolbox AMICI (Fröhlich et al., 2017b), which provides a MATLAB, python and C++ interface to the ODE solver CVODES (Serban and Hindmarsh, 2005) from the SUNDIALS package. As Newton’s method was not applicable due to singular Jacobians, we refactored the code for identifying the steady state from the original publication, which integrates the ODE until the maximum norm of the right hand side undergoes a threshold. Using AMICI, we achieved an average speed-up of about a 200-fold when compared with the original MATLAB implementation (Figure 2A). As the stoichiometric matrix of the model contained non-integer entries, slightly negative states caused problems when exponentiated with these non-integer numbers. Hence, we tested the influence on numerical stability of either adapting the stoichiometrix matrix by rounding these entries – which results only in an approximation of the original model – or enforcing positivity of the states, by setting negative states to zero, by running model simulation and gradient evaluation for 500 randomly sampled parameters (Figure 2B). Particularly for gradient computation, reliability was markedly improved. Yet, likelihood computation often failed, if no steady state could be identified for the given parameter vector.

**Figure 1:**
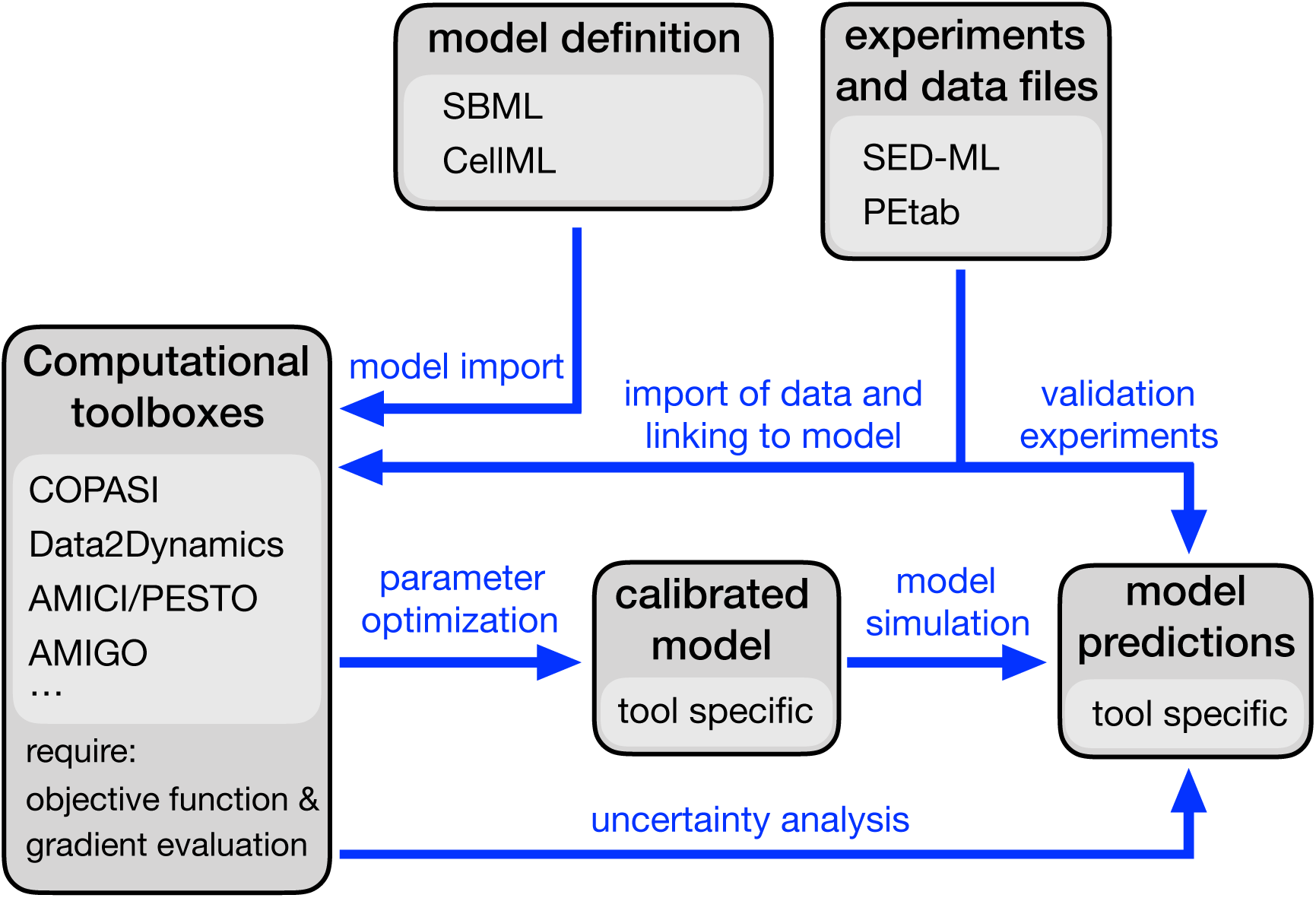
Schematic overview how a scientific question can be addressed using a parametric model. Boxes indicate model, data and tools, arrows indicate actions. Ideally, for all of them standards or best-practices are available, which saves time and reduces errors.

**Figure 2:**
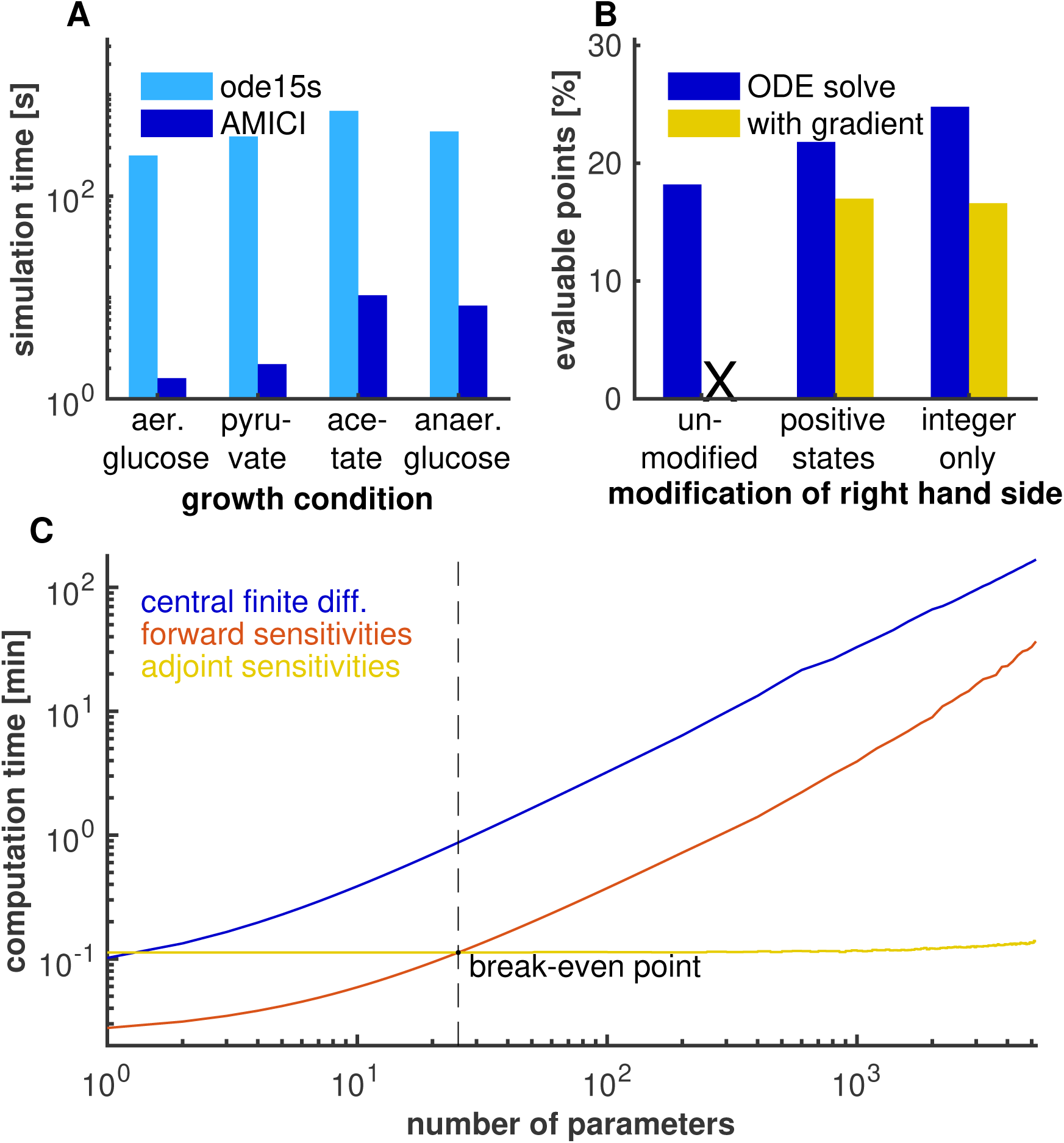
Analysis of model simulation. A) Comparison of computation times using the MATLAB based ODE solver `ode15s` and the solver package AMICI, called from MATLAB but based on C/C++ code (for simulation condition taken from the original publication). B) Comparison of ODE solver robustness for the original model implementation and two modifications (rounded entries in the stoichiometric matrix and enforced positivity of states) for 500 randomly sampled parameter vectors in vicinity of the reported parameter vector. C) Scaling behaviour of computation time for the objective function gradient (one experimental condition) using either finite differences, forward sensitivities or adjoint sensitivities after fixing different numbers of parameters.

Generally, achieving an efficient and robust model simulation is a major issue. Using mature toolboxes with customizable settings and solver diagnostics may help to reduce computation time and increase the robustness.

### 3.4 Objective function and gradient computation

Given observables and measurement data, an objective function can be computed and optimized. As many optimization algorithms require the gradient of the objective function to work, their efficiency depends on the accuracy and the computation time of the objective function gradient. For large-scale models, gradient calculation is a major challenge, as it is computationally substantially more expensive than a usual model simulation.

Standard methods, such as finite differences and forward sensitivities, scale with the product of the dimensions of the state vector and the parameter vector, which makes them computationally prohibitive for large-scale models. This problem can be solved by using adjoint sensitivity analysis, which only needs to solve the original ODE, the ODE of the adjoint state (which has the same size), plus one-dimensional quadratures for each model parameter. For large-scale models, this can reduce the computational burden by two or three orders of magnitude (Fröhlich et al., 2017a). If only steady state data is considered, computing steady state sensitivities can be an alternative. Those can be obtained by directly differentiating Equation (3) with respect to the model parameters, which yields 0 = ∇_*x*_*f* ∇_*θ*_*x* + *∂*_*θ*_*f*, and factorizing the Jacobian of *f*. This approach has the potential to even outperform adjoint sensitivity analysis, but it is only applicable if the Jacobian can be factorized.

In our application example, the Jacobian was singular and the threshold we used for the norm of the right-hand side was too rough to rely on the approach via steady state sensitivities. Following the original implementation, we first identified a time point *t*^*∗*^ for the steady state *x*^*∗*^ for each experimental condition by integrating the ODE until the right hand side became small. Then, we used the experimental data together with the time point *t*^*∗*^ for adjoint sensitivity analysis. We compared the computation time needed to calculate a gradient of the objective function at the nominal parameter value (Figure 2C). We found that adjoint sensitivity analysis outperformed forward sensitivity analysis by a factor of more than 250 and was 1200 times faster than using finite differences. However, the adjoint method only worked reliably after we ensured positivity of the entries in the state vector or rounded the entries in the stoichiometric matrix to integer values (Figure 2B).

If gradients are needed for parameter estimation of a large-scale model, adjoint sensitivity analysis is currently the most efficient method. If only steady state data is used, steady state sensitivities may also be considered, as those do not only yield the gradient, but even state sensitivities. The latter allow the computation of the FIM, which can be valuable for either specific optimization or uncertainty analysis strategies.

### 3.5 Parameter optimization

Parameter estimation for ODE models is a difficult task, as those often yield non-convex and multi-modal objective functions. However, for large-scale models it becomes even more challenging, since the parameter space is high dimensional and therefore harder to explore, and since the objective function and its gradient are more expensive to compute. Yet, ODE models need calibration to produce meaningful outputs.

It has shown to be efficient to exploit as much prior knowledge about the problem structure as possible. Among others, estimating parameters in logarithmic scale (Hass et al., 2019) generally improves the convexity of the objective function and optimizer performance. Moreover, due to the specific form of the objective function, observation specific parameters such as scaling or measurement noise parameters can be computed analytically for a given model simulation, which also increases optimizer performance (Loos et al., 2018; Schmiester et al., 2019). In comparative studies of optimization strategies, meta-heuristics combined with local searches or naive multi-start local optimization have shown to be well suited for this class of problems (Raue et al., 2013; Villaverde et al., 2018). Concerning sub-strategies for local optimization, those using derivative information and hence exploiting the smoothness of the problem have shown to outperform derivative-free methods (Schälte et al., 2018).

Khodayari and Maranas (2017) found their reported parameter vector by a particularly tailored genetic algorithm. We used this vector and sampled parameters in a box of *±*3 around it after a log_10_-transformation. Starting from these initial guesses, we tested different optimization strategies: a multi-start local optimization. with 500 starts using for local optimization either the gradient-based interior-point algorithm of the MATLAB routine `fmincon`, a MATLAB implementation of the gradient-free Dynamic Hill Climbing (De La Maza and Yuret, 1994), and 10 starts of the global, gradient-free genetic algorithm from the toolbox MEIGO (Egea et al., 2014) without local searches (which could however have been combined with gradient-based methods). To our surprise, none of the employed methods was able to find a parameter vector which gave an equally good fit as the reported parameters, although the gradient-based algorithm `fmincon` came closest to it (Figure 3). Admittedly, it could not improve the fit when started at the nominal parameters, in contrast to DHC, which however turned out inferior to `fmincon` when started at randomly sampled guesses. Finally, the purely derivative-free genetic algorithm provided the poorest fit.

**Figure 3:**
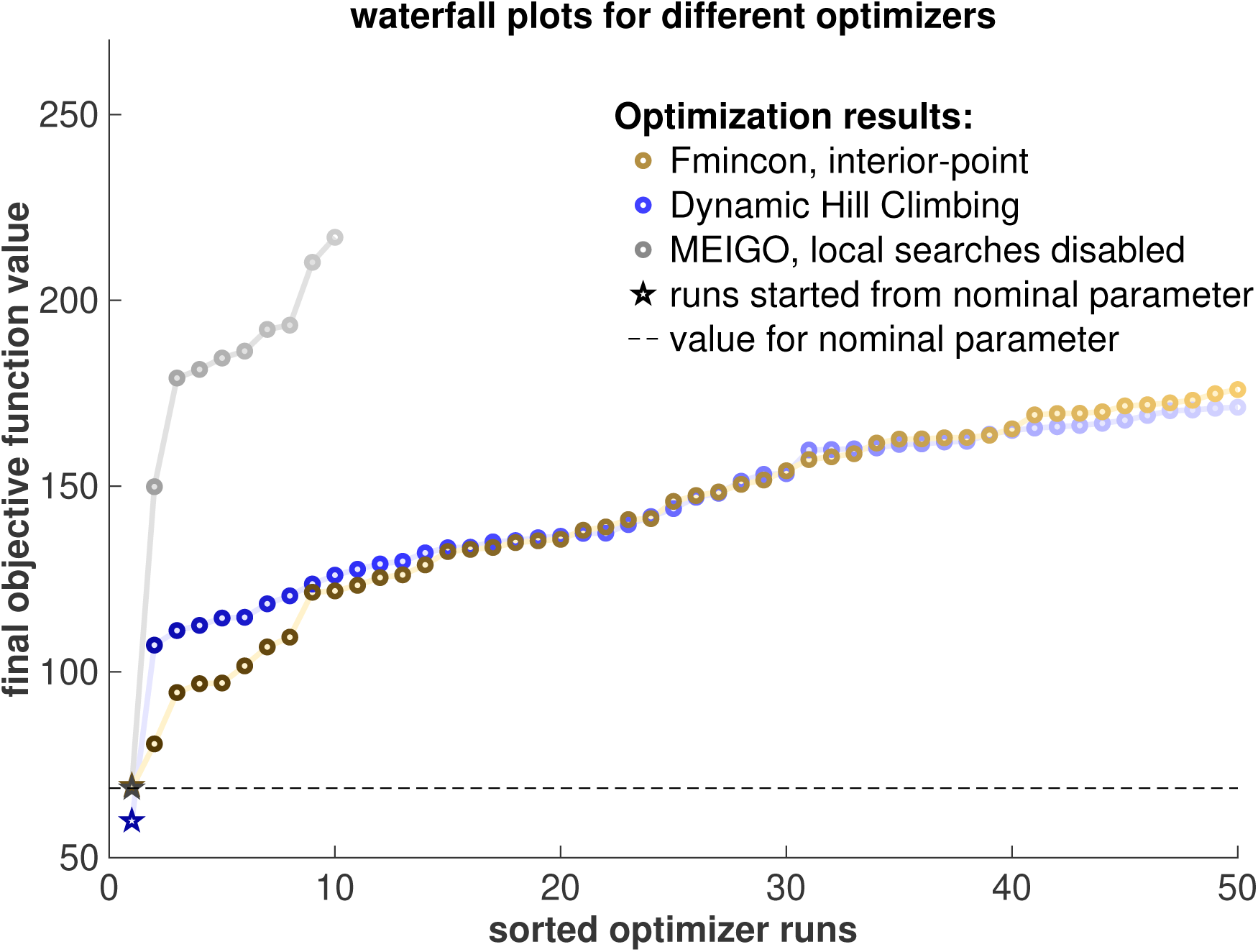
Waterfall plot showing results of different optimizers using multi-start optimization. For the local optimizers fmincon and DHC, the best 50 starts are depicted. The toolbox MEIGO was used as global optimizer, only the gradient-free genetic algorithm was used and therefore started less often.

Overall, this shows that parameter estimation methods which are tailored to the problem tend to perform best. Apart from this, it confirms the known finding that derivative-based methods often outperform derivative-free methods on smooth objective functions, especially if gradients can be computed efficiently. Despite the use of accurate gradients, parameter optimization tends to become harder the larger the considered model is.

### 3.6 Uncertainty analysis

Large-scale models possess many degrees of freedom and therefore, parameters are often underdetermined. Badly determined (i.e., non-identifiable) parameters can (but do not need to) lead to large uncertainties in model predictions. As uncertainty analysis is computationally highly demanding, this is probably the greatest challenge to date which has to be faced when working with large-scale models. Various methods exist in this field, and generally, the methods which yield more insight, such as profile likelihoods or Markov-chain Monte Carlo (MCMC) sampling, are computationally more expensive. Profile likelihoods have to be computed for each parameter (which yields a linear scaling) and for every profile, many (local) optimization problems have to be solved, which in turn are more expensive the larger the model is. MCMC sampling methods try to explore the whole parameter space, which means more function evaluations are needed to create a representative sample in high dimensions. Generally, their scaling behavior is hard to assess, although for particular posterior distributions estimates can be derived, which generally suggest a polynomial scaling in the parameter dimension. Therefore, profile likelihoods and MCMC sampling often become prohibitive for large-scale models.

Different approaches for this problem exist: For profile computation, cheaper methods have been developed, which try to circumvent optimization (Stapor et al., 2018) by integrating a continuous analogue of the profile optimality condition. Furthermore, profiles can be computed directly for model predictions (Kreutz et al., 2012), which may be beneficial if model predictions are less numerous than model parameters. Yet, if parameter optimization does not work reliably, these methods are not applicable, as they need an optimization result for initialization. Consequently, cheaper and less accurate methods are mostly used, such as evaluating the variance in ensemble predictions (Henriques et al., 2017), analyzing the variance of the optimized parameters or gradients, using local approximations such as the Fisher information matrix (FIM), or splitting the data in different training and validation sets and applying cross-validation (Froehlich et al., 2018), if the data set is large enough.

For our application example, profile likelihoods and MCMC sampling were computationally too expensive. Hence, we computed the FIM at the reported parameter value based on forward sensitivity analysis (see e.g. Raue et al. (2013)) and computed its spectral decomposition. After rescaling the matrix with the inverse of the largest eigenvalue, we set a cut-off at 10^*-*15^, which corresponds roughly to the numerical precision of MATLAB. This yielded 675 identifiable (and hence 4564 non-identifiable) directions in parameter space (Figure 4).

**Figure 4:**
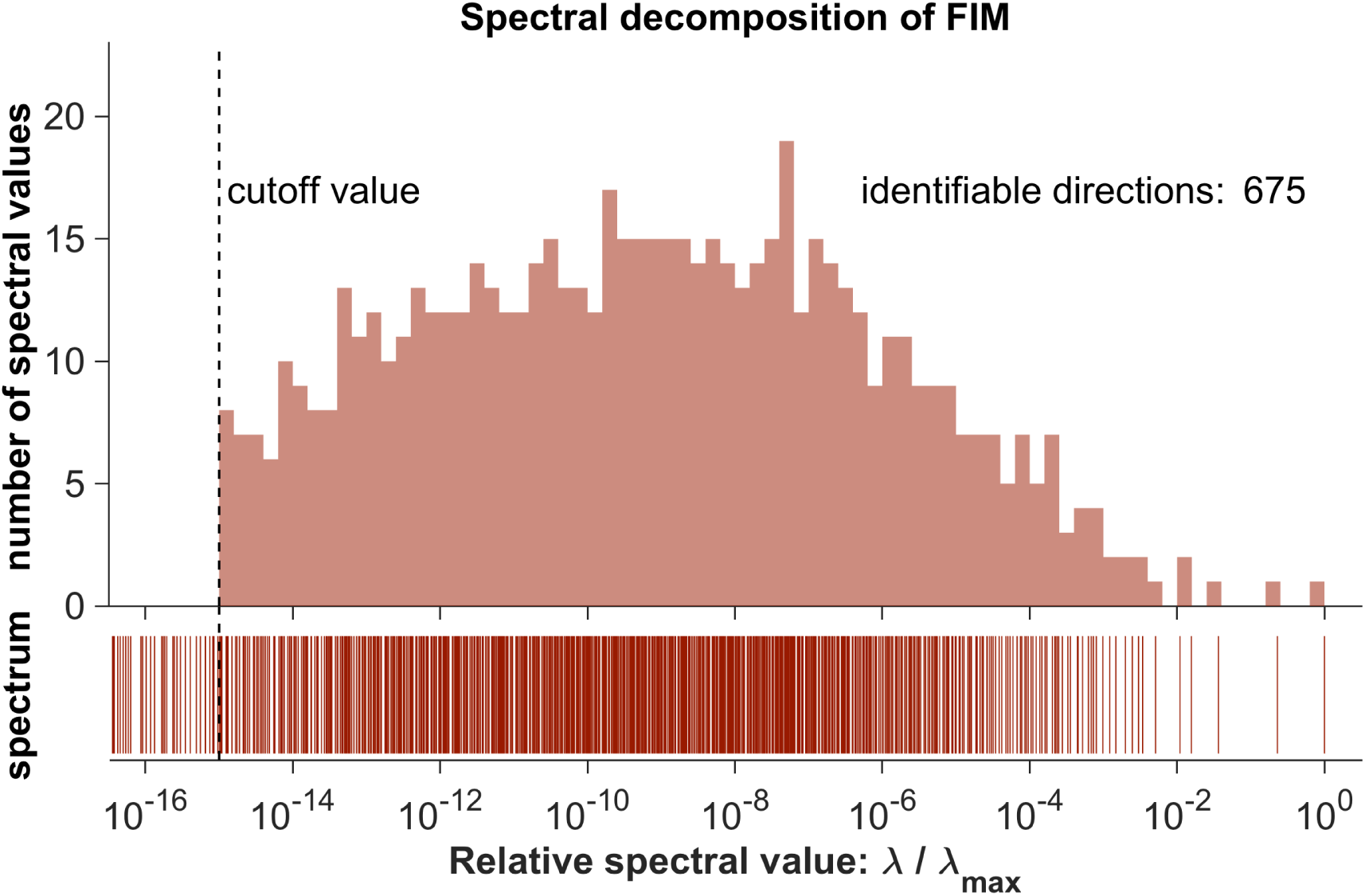
Spectral decomposition of the FIM. Vertical ticks indicate spectral values, the histogram shows their distribution.

Overall, uncertainty analysis is probably the most challenging task for large-scale models. Mainly simple approximations are available, which is a highly problematic situation, as large-scale models are expected to suffer more from uncertainties than smaller-scale models. Probably, uncertainties of predictions should rather be inferred directly than going via parameter uncertainties. For this purpose, splitting the available data into different combinations of training and validation sets and performing cross validation is probably the most practicable method at the moment.

## 4 Summary and conclusion

Large-scale modelling with ODEs is a rapidly growing field, which comes with new challenges. In this review, we summarized the most recurrent problems and showed currently available approaches to deal with them, which we illustrated on a recent application example. For many problems, no general solutions exist. More models with measurement data, collected in public data bases using standards, are needed to perform further studies.

Leaving aside the even more complicated task of network inference, parameter optimization and uncertainty analysis are currently the key challenges for large-scale models, for which no satisfactory approaches exist. Due to the high computation times, toolboxes suitable for computing clusters are necessary and have recently been developed (Penas et al., 2017; Schmiester et al., 2019). Moreover, new approaches have to be explored, such as transferring the concept of mini batching from the field of deep learning to optimization (Goodfellow et al., 2016) or MCMC sampling (Seita et al., 2017) of dynamic models, and must be adapted to ODE models.

As methods which are tailored a problem class tend to outperform black box solutions and since parameter estimation of ODE models is a bounded field, accounting for the specific structure can lead to substantial improvements (Froehlich et al., 2018; Schmiester et al., 2019). Studies have to be performed, which aim at a better understanding of the properties of this problem class, such as how non-identifiable parameters translate into uncertainties of model predictions.

First steps have been taken to facilitate the study of this new field. Now, we have to deepen our understanding of the problems at hand. Hopefully, this will enable the development of novel strategies to tackle the existing challenges and lead to a substantially improved understanding of complex biological questions in the future.

## Acknowlegdements

We want acknowledge the help and advise of Fabian Fröhlich for finding errors and spotting origins of numerical issues in our implementation of the application example, as well as the help of Prof. Maranas and his work group when answering the questions we had concerning their model and their implementation.

This work was supported through the European Union’s Horizon 2020 research and innovation programme under grant agreement no. 686282 (J.H., P.S.).

